# Domestication signatures in the non-conventional yeast *Lachancea cidri*

**DOI:** 10.1101/2023.06.12.544598

**Authors:** Pablo Villarreal, Samuel O’Donnell, Nicolas Agier, Felipe Muñoz-Guzmán, José Benavides-Parra, Kamila Urbina, Tomas A. Peña, Mark Solomon, Roberto F. Nespolo, Gilles Fischer, Cristian Varela, Francisco A. Cubillos

## Abstract

Evaluating domestication signatures beyond model organisms is essential for thoroughly understanding the genotype-phenotype relationship in wild and human-related environments. Structural variations (SVs) can significantly impact phenotypes playing an important role in the physiological adaptation of species to different niches, including during domestication. A detailed characterization of the fitness consequences of these genomic rearrangements, however, is still limited in non-model systems, largely due to the paucity of direct comparisons between domesticated and wild isolates. Here, we used a combination of sequencing strategies to explore major genomic rearrangements in a *Lachancea cidri* yeast strain isolated from cider (CBS2950) and compared them to those in eight wild isolates from primary forests. Genomic analysis revealed dozens of SVs, including a large reciprocal translocation (∼16 kb and 500 kb) present in the cider strain, but absent from all wild strains. Interestingly, the number of SVs was higher relative to single-nucleotide polymorphisms in the cider strain, suggesting a significant role on the strain’s phenotypic variation. The set of SVs identified directly impacts dozens of genes, and likely underpins the greater fermentation performance in the *L. cidri* CBS2950. Additionally, the large reciprocal translocation affects a proline permease (*PUT4*) regulatory region, resulting in higher *PUT4* transcript levels, which agrees with higher ethanol tolerance, improved cell growth when using proline, and higher amino acid consumption during fermentation. These results suggest that SVs are responsible for the rapid physiological adaptation of yeast to an anthropogenic habitat and demonstrate the key contribution of SVs in adaptive fermentative traits in non-model species.

**Importance:** The exploration of domestication signatures associated with anthropogenic niches has predominantly focused on studies conducted on model organisms, such as *Saccharomyces cerevisiae*, overlooking the potential for comparisons across other non-*Saccharomyces* species. In our research, employing a combination of long– and short-read data, we found domestication signatures in *L. cidri*, a non-model species recently isolated from fermentative environments in cider in France. The significance of our study lies in the identification of large array of major genomic rearrangements in a cider strain compared to wild isolates, which underly several fermentative traits. These domestication hallmarks result from structural variants, which are likely responsible for the phenotypic differences between strains, providing a rapid path of adaptation to human-related environments.

## Introduction

Domestication is an evolutionary process resulting from a specialized mutualism, in which one species controls the fitness of another to gain resources or services (1). Human-associated domestication, such as for plants and animals, is undoubtedly a well-known process in human history (1, 2). Interestingly, while the domestication of some developmentally complex organisms is the result of intentional human behavior, this process happened largely unintentionally in microorganisms (1, 3). The constant interaction of microorganisms in human-related processes allowed their transition from variable and heterogeneous environments, typical of wild ecosystems, to more stable and predictive ones, characteristic of anthropogenic niches, over time (4).

Genomic variation in a population in response to environmental changes is shaped by diverse biological processes, including genetic drift, natural selection, and migration (5, 6). Historically, most studies on genetic variation have focused on single-nucleotide polymorphisms (SNPs), as high-throughput sequencing technologies provide massive amounts of information for comparing species, populations, calibrating divergence times, and inferring adaptive traits of interest (7–12). Recent studies in several organisms, however, have revealed that structural variations (SVs) can sometimes better explain the observed phenotypic diversity of populations (13, 14). Indeed, experimental evolution assays have indicated that SVs are an important driver of evolution and adaptation to new environments, such as to human-related niches (14). SVs are defined as a region of DNA that shows a change in copy number (CNV), orientation, or chromosomal location between individuals (12). SVs can be balanced and have no specific loss or gain of DNA information (inversions and translocations), or they can be unbalanced, where a fraction of the genome is lost or duplicated (insertions, deletions, and duplications) (7, 12).

There are numerous examples in fungal species of physiological adaptations to anthropogenic niches derived from SVs (13, 15–19). For example, wine, beer, and cider represent three anthropogenic environments where different microorganisms are responsible for the fermentation process, including one of the best-studied domesticated microorganisms, the yeast *Saccharomyces cerevisiae*. In wine, greater sulfite resistance in different species across the *Saccharomyces* genus emerged because of major genomic rearrangements, including inversions and chromosomal translocations, impacting the expression of the *SSU1* gene, a sulfite efflux pump (13, 17, 20–22). These studies demonstrated that different genomic variations associated to SVs could be responsible for the rapid adaptation of microorganisms to human-related environments. However, whether there are additional examples of genomic variation events associated to fermentation-related domestication is still largely unknown, particularly in organisms other than *S. cerevisiae.* A better understanding of these events and their consequences can guide biotechnological solutions to improve these industrial processes.

Recent bioprospecting studies in human environments, like wine and cheese, have expanded the repertoire of domesticated yeast strains (23). For instance, genomic changes associated to anthropogenic habitats have been reported in different non-conventional yeast genera, such as *Kluyveromyces* and *Torulaspora* (8, 19, 24–26). In all, these studies have deepened our understanding of fungi adaptation to anthropogenic niches, however, additional detailed molecular evidence is needed to understand the adaptative process behind human-made environments and the role of genomic plasticity underlying such adaptation. Cider is a complex fermentative environment where dozens of species interact and sequentially develop throughout the process (27). One of the predominant yeasts in cider fermentation is *Lachancea cidri* (28). This species diverged over 150 MYA from *S. cerevisiae*, it lacks a known sexual cycle, and only haploid strains have been so far recovered (29, 30). This yeast has been isolated from cider fermentation environments in France (reference strain CBS2950), and primary forests in Australia and Patagonia, exhibiting different genetic and phenotypic patterns depending on the isolation environment (30, 31). Despite the diverse ecosystems where *L. cidri* strains are present, there is little information about their adaptation to anthropogenic niches at the genetic level.

To study the physiological adaptation of a non-conventional yeast to a human-related environment, specifically, cider fermentation, we explored the genetic landscape of *L. cidri* under different fermentative conditions. We generated telomere-to-telomere genome assemblies of wild strains to identify key genomic signatures underlying phenotypic differences. We focused on SVs and explored how these might have impacted fermentative capacity and could have ultimately allowed a shift from a wild lifestyle to an anthropized one. Overall, our study offers new insights into the evolutionary history of a yeast species and explains how human-related environments can prompt genome reorganization as a hallmark of domestication.

## Results

### Anthropogenic *Lachancea cidri* strain exhibits phenotypic differences under fermentative conditions compared to wild strains

To explore phenotypic differences between *L. cidri* anthropogenic and wild strains, two fermentation assays were performed in natural musts: apple juice, to produce cider, and Chardonnay grape must, to produce wine. For both assays, three strains were considered: 1) the domesticated strain *L. cidri* CBS2950, obtained from a cider fermentative environment in France, 2) the wild LC1 strain, which is the closest known relative to CBS2950 and was isolated from tree sap samples in the Central Plateau of Tasmania, Australia, and 3) NS18, a wild *L. cidri* strain from *Nothofagus* forests from Patagonia (Table S1) (30, 32).

No significant differences in fermentation kinetics were observed among the *L. cidri* strains during cider fermentation. The three *L. cidri* strains showed a similar CO_2_ loss (g/L) profile (*p-value* > 0.05, one-way ANOVA) compared to the *S. cerevisiae* Lalvin EC1118^®^ commercial control strain (*p-value* > 0.05, one-way ANOVA), demonstrating their remarkable ability to ferment this apple must (Fig S1A). Furthermore, while no differences were observed among the *L. cidri* strains in the production of various metabolites, such as ethanol, glycerol, and succinic acid (Fig S1, *p-value* > 0.05, one-way ANOVA), in the case of ethanol and succinic acid their production was higher in these strains than in the control (Fig S1B-D *p-value* < 0.05, one-way ANOVA).

In Chardonnay grape must, the three *L. cidri* strains completed the fermentation after 15 days, while the *S. cerevisiae* control strain finished in 4 days (Fig 1A). Yet, at the end of the Chardonnay fermentation, ethanol concentration was significantly higher in the cider strain compared to that in all of the other strains, including the *S. cerevisiae* control (*p-value* < 0.05, one-way ANOVA) (Fig 1B). While no significant differences among the *L. cidri* strains were observed to produce glycerol and succinic, citric, lactic, and acetic acids (Fig S2, *p-value* > 0.05, one-way ANOVA), the *S. cerevisiae* control strain produced higher succinic and citric acid concentrations than all *L. cidri* strains (Fig S2, *p-value* > 0.05, one-way ANOVA). A particular case was the production of malic acid, where the lowest production was observed in strain NS18 (Fig S2, *p-value* < 0.05, one-way ANOVA), while CBS2950 and LC1 did not show significant differences between them (Fig S2, *p-value* > 0.05, one-way ANOVA).

**Figure 1.**
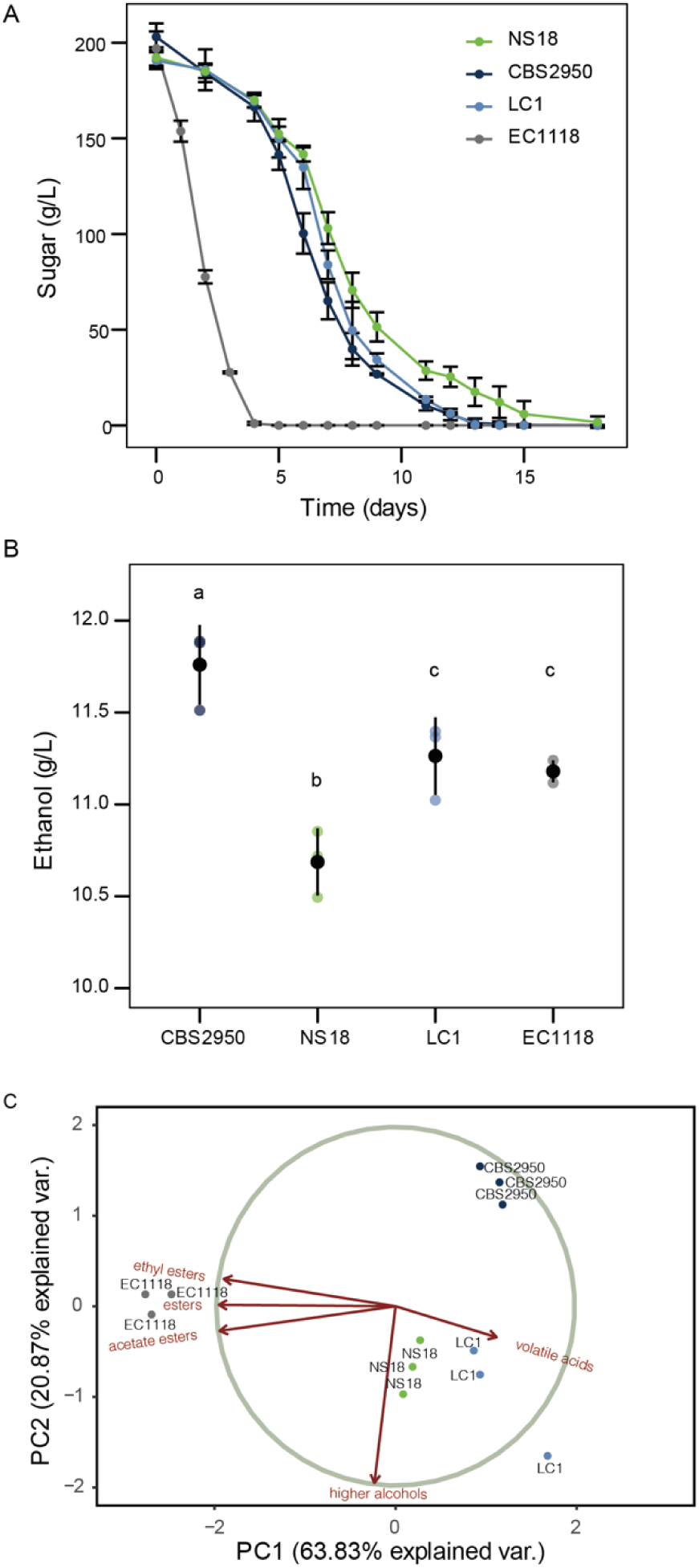
*Lachancea cidri* fermentation performance under Chardonnay wine must. (A) Kinetics of sugar consumption [g/L] in Chardonnay fermentations (n=3). (B) Ethanol production [g/L] at the end of the Chardonnay fermentation. (C) Principal Component Analysis of Volatile Compound production in Chardonnay fermentation. Different letters reflect statistically differences between strains with a *p*-value < 0.05, one-way analysis of variance (ANOVA).

Distinctive production profiles of volatile compounds (VCs) were observed for strains isolated from the fermentative environments, *L. cidri* CBS2950 and *S. cerevisiae* Lalvin EC1118^®^, compared to wild strains (Table S3). To reduce the dimensionality of the dataset and interpret the VC production for each strain, a global principal component analysis (PCA) using the VCs data was performed (Fig 1C). The first two components explained 63.8 % and 20.9 % of the observed variation, respectively separating the strains according to their isolation source (forests, wine, or cider) (Fig 1C). The CBS2950 strain produced lower concentrations of higher alcohols and acetate esters (*p-value* < 0.05, one-way ANOVA, Table S2) than the other *L. cidri* strains, demonstrating a distinctive organoleptic profile and revealing a clear phenotypic differentiation from the wild strains.

Altogether, these results highlight phenotypic differences in the CBS2950 strain compared to wild *L. cidri* strains, suggesting a specific adaptation to the fermentative environment from its original isolation source.

### *Lachancea cidri* cider strain features major chromosomal rearrangements compared to wild strains

The CBS2950 and LC1 *L. cidri* strains differ by only 41 SNPs across the genome, and none of these appear to impact protein-coding sequences (30). To identify the genetic basis of the phenotypic differences between these two strains, we decided to explore genetic changes beyond SNPs and to evaluate the presence of SVs. A molecular karyotype using these two strains plus eight wild strains from different localities was first performed to identify major rearrangements in the genome, using pulse-field electrophoresis (Fig 2A, Table S1). The chromosomal DNA of the different strains was separated into 8 electrophoretic bands ranging in size from 300 to 2,700 kb (Fig 2A). Chromosomes LACI0B and LACI0G exhibited a unique migration pattern in CBS2950, with significant size variation compared to the other strains (Fig 2A).

**Figure 2.**
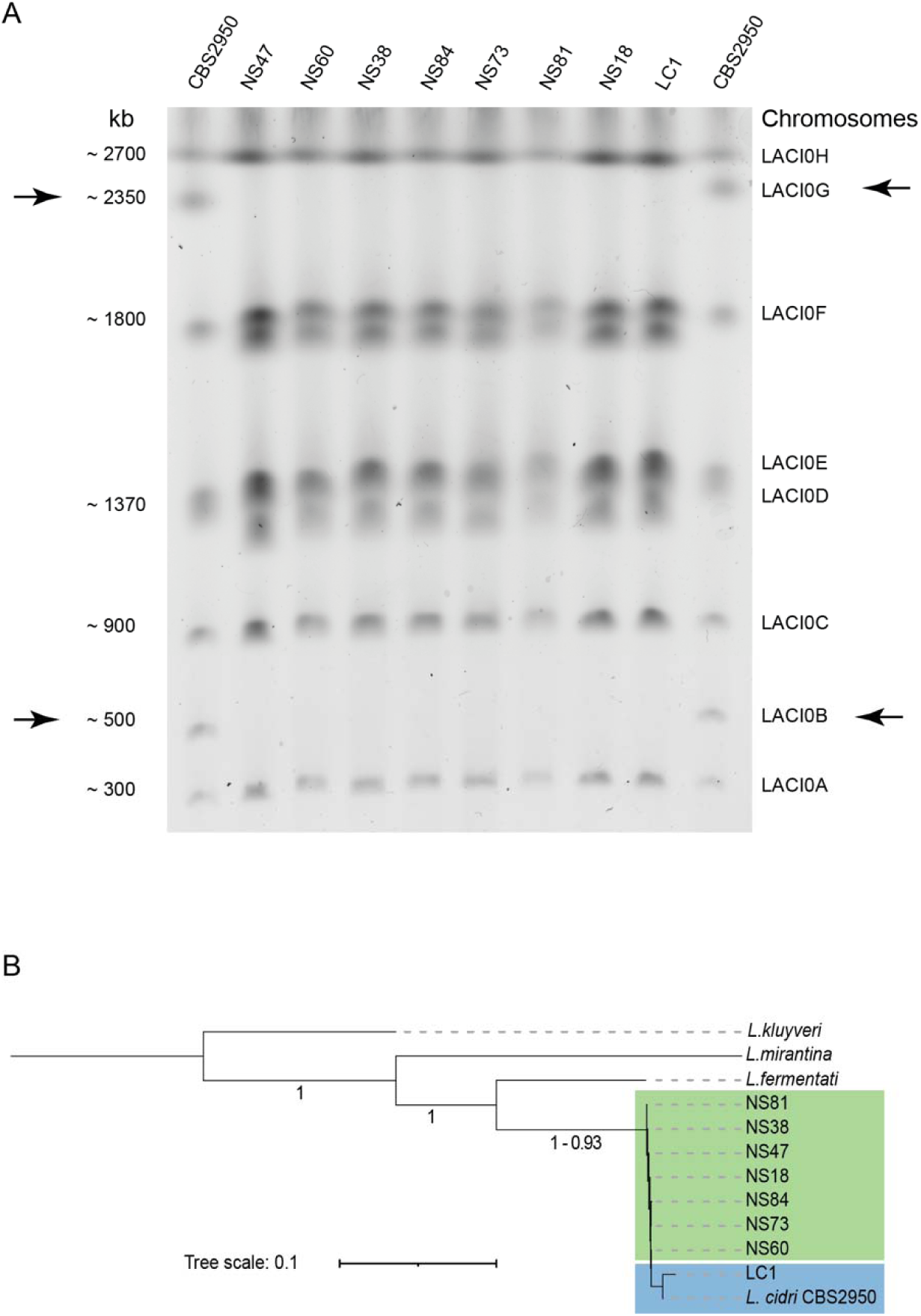
Molecular karyotype and phylogeny of *L. cidri* strains. (A) Pulsed-field electrophoresis of the chromosomal DNA of *L. cidri* strains. *L. cidri* CBS2950 is located in the first and last column. Wild South American strains: NS47, NS60, NS38, NS84, NS73, NS81, NS18. Wild Australian strain: LC1. Black arrows show the chromosomes with a different electrophoretic pattern between wild and cider strain. (B) Consensus phylogenetic tree of *L. cidri* long-read genomes. The tree was built by orthogroup inference. Branch lengths represent the average number of substitutions per site across the sampled gene families. *L. mirantina*, *L. kluyveri,* and *L. fermentati* were used as outgroups.

Whole-genome long-read Nanopore sequencing was used to assemble the genome of the eight wild *L. cidri* strains (one contig per chromosome) in an attempt to identify the origin of the chromosome size differences (Fig S3, Table S3, and S4). Using a consensus species tree, we observed clustering between *L. cidri* CBS2950 and *L. cidri* LC1 (Fig 2B), confirming the high genetic relatedness between these two strains, consistent with previous studies (30). Strains isolated from wild environments are at the base of the species tree, while *L. cidri* CBS2950 recently diverged from the Australian clade.

We then evaluated the presence of structural variants (SVs) in these strains by comparing all *de novo* assemblies against the *L. cidri* CBS2950 strain. Perfect collinearity was observed in all chromosomes, except for chromosomes LACI0B and LACI0G in the CBS2950 strain (Fig 3A). This strain showed a ∼16-Kb and 500-Kb terminal translocation between chromosomes LACI0B and LACI0G, respectively (Fig 3A), which is consistent with the karyotyping results (Fig 2A). Comparing two wild strains randomly, perfect collinearity was observed across all chromosomes, demonstrating that chromosome size differences are specific to the cider strain (Fig 3B). To quantify the extent of all SVs across the genomes, a comprehensive analysis using pairwise comparisons (MUM&Co) was performed (Fig 3C). Six types of SVs were evaluated: deletions, insertions, duplications, contractions, inversions, and translocations (Table S5). We found that the cider strain had a higher amount of total variation (mean = 71.1 SV count) compared to the closely related wild strain *L. cidri* LC1 (mean = 52.8 SV count) (Fig S4A), demonstrating enrichment of SVs relative to SNPs in CBS2950 across the genome (Fig 3D). Most of the structural variants were localized in sub-telomeric regions (Fig S4B), mainly corresponding to unbalanced variants (Fig S5C). Overall, the CBS2950 strain showed a higher number and larger size of SVs than the other wild strains (Fig S5A and B), with the most frequent SV being deletions (Fig S6A). In addition, we found in CBS2950 a higher frequency of SVs impacting intergenic regions compared to coding sequences (Fig S6B), which may affect gene expression.

**Figure 3.**
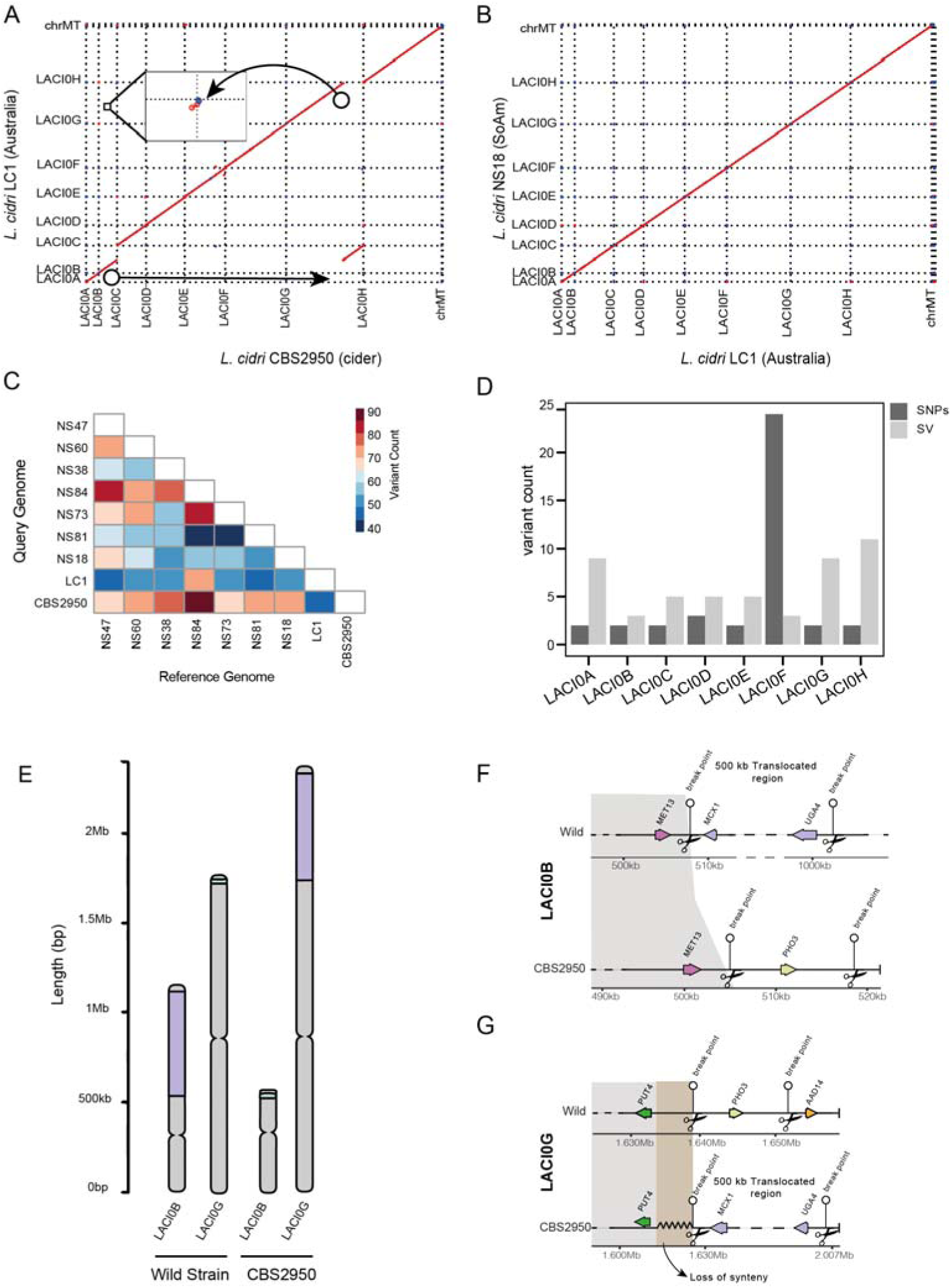
*L. cidri* long-read genome assembly. (A) Genome synteny analysis of the reference strain CBS2950, and the Australian LC1 strain. (B) Genome synteny analysis of two wild *L. cidri* strains LC1 and NS18. Dot plot representation of DNA sequence identity. Red dots indicate forward matches and blue dots for reverse matches. (C) SVs pairwise comparisons among all *L. cidri* genome assemblies with the total number of structural variations. (D) LACI0B and LACI0G chromosome comparison between wild and CBS2950 strains. Purple and green colors denote translocated regions. (E) Number of SVs and SNPs per chromosome between CBS2950 and LC1 (F) LACIB Chromosome synteny (G) LACIG Chromosome synteny. Black scissors show breakpoints in the genome.

To compare the impact of SV on gene order, we performed a genome comparison between strains across the genome. A perfect synteny was observed between the CBS2950 strain and the wild strains, except for the LACI0GtLACI0B translocation (Fig 3A and 3E). This translocation comprises 226 genes, which are relocated from chromosome LACI0B to chromosome LACI0G in the CBS2950 strain. The translocation breakpoint also impacted the regulatory region of the *PUT4* gene in the cider strain, which encodes for a proline permease required for high-affinity proline transport. These genomic data demonstrate a large genetic remodeling in the domesticated *L. cidri* CBS2950 strain compared to its wild counterparts.

### *Lachancea cidri* isolates exhibit different fermentation capacity under varying nitrogen concentrations

To evaluate whether phenotypic differences between wild and anthropogenic *L. cidri* strains are related to nitrogen consumption, we performed micro-fermentation assays under different nitrogen conditions, comparing CBS2950 and the LC1 strain. For this, micro-fermentations using Synthetic Wine Musts (SWM) with different yeast assimilable nitrogen (YAN) concentrations, SWM300 (300 mg/mL YAN) and SWM60 (60 mg/mL YAN), were performed. In SWM300, no differences were observed at the end of the fermentation between these *L. cidri* strains regarding fermentation kinetics or total CO_2_ production (*p-value* > 0.05, one-way ANOVA) (Fig 4A). Conversely, a decrease in the amount of YAN (SWM60) had a significant impact on the fermentation performance, and the extent of the effect depended on the strain (*p-value* < 0.05, one-way ANOVA, Fig 4A). Under low-nitrogen conditions, the CBS2950 cider strain exhibited a higher fermentative capacity, suggesting a greater ability to ferment under low-nitrogen conditions (Fig 4A). The CBS2950 strain exhibited a higher and more efficient nitrogen utilization profile than the wild strain LC1 early in the fermentation under both conditions (*p-value* < 0.05, one-way ANOVA, Fig 4B and C, Table S6). However, such more efficient nitrogen consumption in SWM300 did not necessarily resulted in greater fermentation capacity (Fig 4B). This contrasts with the situation in SWM60, where faster nitrogen consumption at earlier time points translated into higher CO_2_ loss levels in the cider strain (Fig 4C).

**Figure 4.**
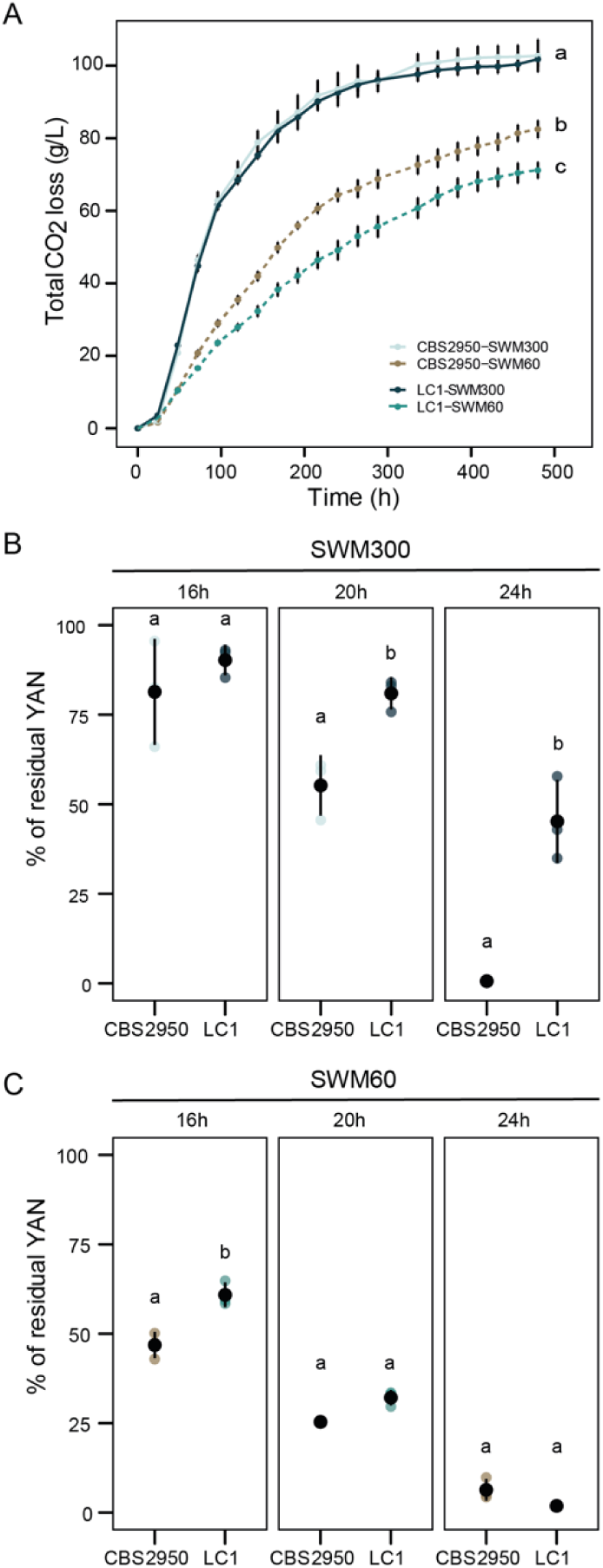
*Lachancea cidri* fermentation performance under Synthetic Wine Must. (A) Fermentation kinetics in Synthetic Wine Must (SWM). The solid line shows the fermentation profiles of the different strains at high nitrogen concentrations SWM300 (300 mg/ml YAN), and the dashed line at low nitrogen concentrations (60 mg/ml YAN). (B) Percentage of residual nitrogen at different time-points in SWM300 fermentation. (C) Percentage of residual nitrogen at different time-points in SWM60 fermentation. Different letters reflect statistically differences between strains with a *p*-value < 0.05, one-way analysis of variance (ANOVA).

To evaluate if fermentation differences impacted the secondary metabolite production, we estimated ethanol, glycerol and organic acids production. No differences were observed in the production of secondary metabolites (ethanol, glycerol, acetic and succinic acid) between the CBS2950 and LC1 in SWM300 (*p-value* > 0.05, one-way ANOVA) (Fig S7). However, in SWM60, differences between CBS2950 and LC1 were observed in the production of glycerol, acetic acid, and succinic acid (*p-value* < 0.05, one-way ANOVA) (Fig S7 C and D). Overall, these results highlight phenotypic differences between cider and wild strains under low nitrogen conditions in synthetic wine must.

### Differential gene expression in the Anthropogenic *Lachancea cidri* strain

We surmised that the various genomic rearrangements identified in the *L. cidri* cider strain may affect the expression of various genes, which may underlie some of the observed phenotypic differences between CSB2950 and LC1 under low nitrogen conditions. To determine the impact of SVs on gene expression in CBS2950 relatively to the closely related LC1 strain, we performed RNA-seq under low-nitrogen wine fermentation conditions (SWM60). Differentially Expressed Gene (DEG) analysis revealed 569 DEGs between the strains (FDR < 0.05, Table S7). Interestingly, several fermentation and nitrogen-related genes were found as DEGs. These included *GNP1* (broad specificity amino acid permease) and *LAP2* (cysteine aminopeptidase with homocysteine-thiolactone activity), both related to amino acid transport. Both genes were up-regulated in the cider strain relative to LC1 (Fig 5A). *ADH4* (alcohol dehydrogenase isoenzyme IV) was also up-regulated in CBS2950 (Fig 5A), consistent with its greater fermentative capacity. Interestingly, expression differences were not enriched in the translocated genes within the LACI0B and LACI0G chromosomes (*p*-value > 0.05 Hypergeometric Test, Total genome P: 0,142; Translocated region P: 0,150), indicating that the translocation did not significantly affect the expression of these genes under these conditions. Enrichment analysis of gene ontology (GO) terms associated with DEGs highlighted the upregulation of ribosomal genes in CBS2950, and the downregulation of secondary metabolite production like organic acids and carboxylic acid (Table S8). Interestingly, these genes were up-regulated in the wild strain (Table S9), which could explain the differences in the VCs profiles found in Chardonnay fermentation, where the wild strain produces a greater repertoire of aromas compared to the cider strain.

**Figure 5.**
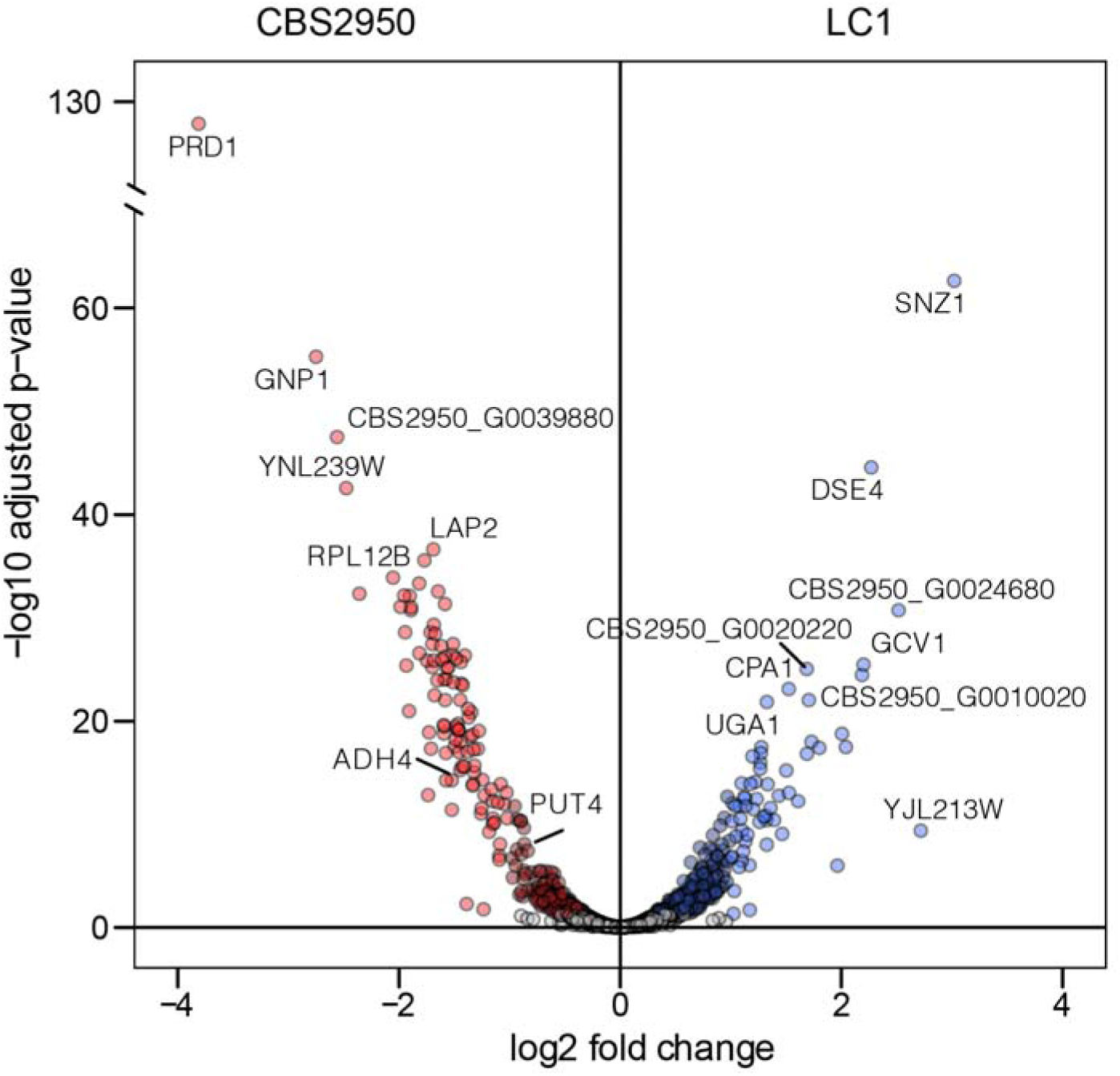
Differential gene expression between wild and anthropogenic *L. cidri* strain under synthetic wine fermentation conditions. A. The volcano plot depicts differentially expressed genes (1-fold change in expression; p-adjusted < 0.05). Up-regulated and down-regulated genes in CBS2950 relative to LC1 are depicted in red and blue dots, respectively.

We also analyzed the region surrounding the LACI0GtLACI0B translocation breakpoints, in the vicinity of the *PUT4* and *PHO3* promoter regions (Fig 3G). While no differences were found for *PHO3, PUT4* was differentially expressed between CBS2950 and LC1 (*p*-adjusted < 0.05, Table S7). In this case, the cider strain exhibited greater *PUT4* expression levels compared to those in LC1, suggesting an effect of the SV on the gene’s expression profile (Table S7). Altogether, these results suggest that changes in the expression of various genes related to nitrogen metabolism may underlie some of the observed phenotypic differences between CSB2950 and LC1 strain under fermentative conditions.

### The Anthropogenic *Lachancea cidri* strain exhibits a series of phenotypic domestication signatures

To determine the presence of domestication signatures in the cider strain, different phenotypic assays representative of the hallmarks of the fermentation process were performed. Given the effect of the translocation event on the expression of *PUT4*, we first evaluated microbial growth using proline as sole nitrogen source (Fig 6A). At each proline concentration assessed, a greater maximum optical density (Fig 6A) and growth rate (μmax) (Fig 6B) were observed in CBS2950 compared to LC1 (*p-value* < 0.05, one-way ANOVA). To determine if this effect is specific to proline, we used a combination of four different amino acids (Glutamic acid, Aspartic acid, Alanine, and Leucine) at 50 and 200 mg/L YAN, and a complete YNB 2% glucose media (Fig S8). In all cases, no significant differences were observed between the two strains (*p-value* > 0.05, one-way ANOVA, Fig S8), suggesting that the growth differences observed are specific to proline.

**Figure 6.**
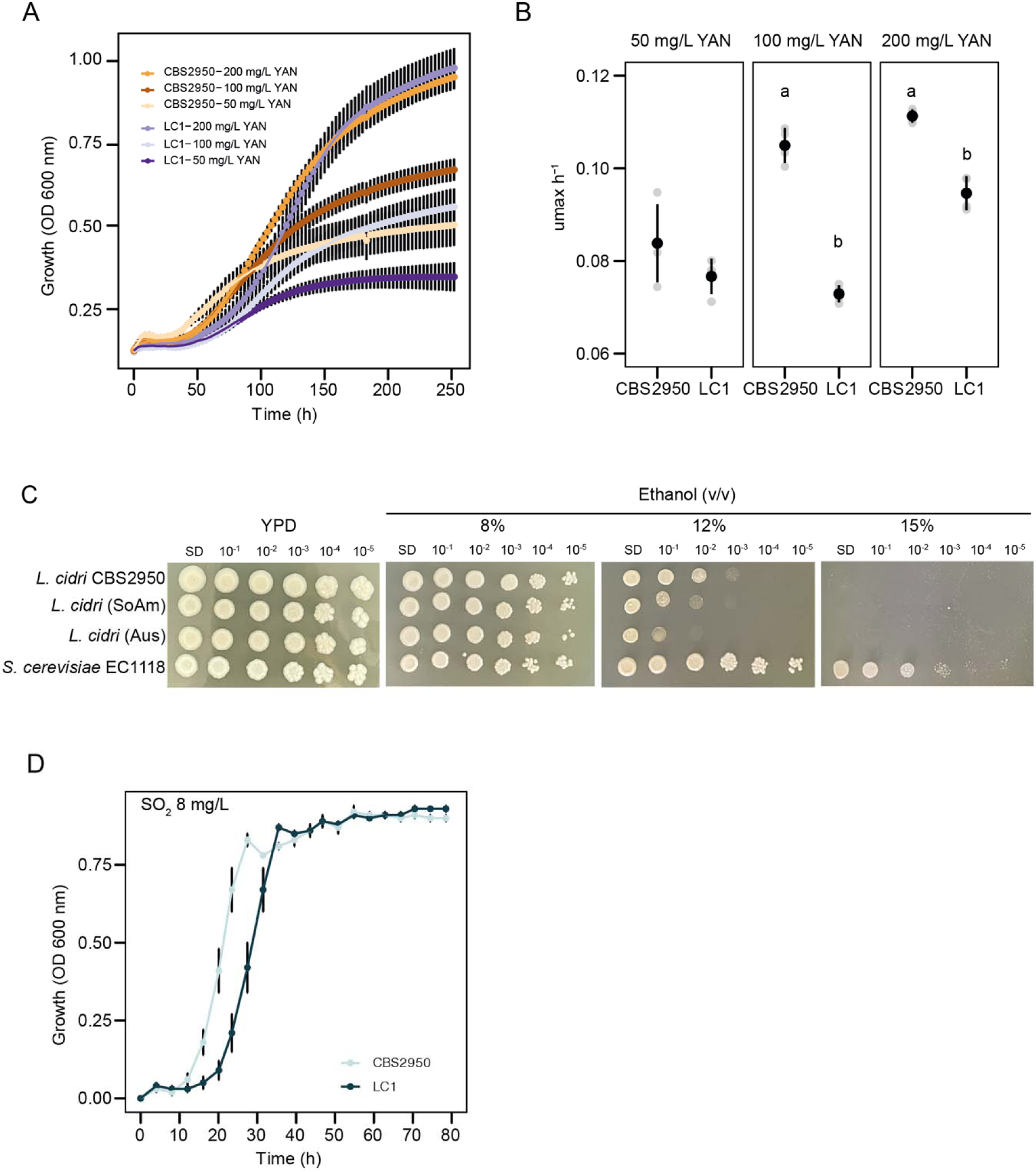
Domestication signatures in *Lachancea cidri*. A. Growth curve at different proline concentrations (mg/L YAN) in the cider (CBS2950) and wild *L. cidri* strains. B. Growth rate at different concentrations of Proline (mg/L YAN). Different letters depict statistically differences between strains with a *p*-value < 0.05, one-way analysis of variance (ANOVA). C. Comparison of ethanol tolerance between the CBS2950 strain and wild (SoAm = NS18, Aus = LC1) strains of *L. cidri*. *S. cerevisiae* EC1118 was used as a control. D. Effect of sulfite (SO_2_ 8 mg/L) on the growth of *L. cidri* strains.

Considering the correlation between the reported function of the *PUT4* gene and ethanol tolerance (33), we also evaluated colony growth under various ethanol concentrations in all *L. cidri* strains (Fig 6C). A similar growth was observed for all strains at 8% v/v, whereas at higher ethanol concentrations (12% v/v) CBS2950 showed improved growth performance, suggesting a greater tolerance and adaptation to high-ethanol concentrations, such as those found in cider fermentation (Fig 6C). In this way, the expression differences in *PUT4* correlated with greater microbial growth when proline was the sole nitrogen source and may be responsible for an increased ethanol tolerance in the “domesticated” *L. cidri* CBS2950 strain compared to the wild *L. cidri* LC1 strain. These results suggest a major role of the SV breakpoint in CBS2950 adaptation to cider environments.

Microbial growth under sulfite (SO_2_) conditions, which is another signature of adaptation to human-related fermentative environments, was also evaluated (Fig 6D and Fig S9). CBS2950 exhibited greater growth performance compared to LC1 under the different SO_2_ concentrations tested (Fig 6D), exhibiting a higher growth rate and a shorter lag phase compared to the native LC1 strain (Fig S9). Given this, we examined the presence of SVs in *SSU1*, a gene that encodes for a sulfite efflux pump known to confer sulfite (SO_2_) resistance (13). Mum&Co and manual analysis of the *SSU1* neighborhood, however, revealed no differences between CBS2950 and LC1.

Altogether, these results demonstrate the presence of different phenotypic advantages under fermentative conditions in the *L. cidri* CBS2950 strain, and these can partly be explained by balanced SVs. This highlights the potential importance of these genomic rearrangements in providing an adaptive advantage to conditions found in anthropogenic fermentative environments.

## Discussion

Domestication is a major selective force characterized by several genomic hallmarks (24, 26, 34, 35). Different forms of genomic reorganization allowing microorganisms to rapidly adapt to novel environmental challenges have been well documented in various model organisms, including yeast and fungi of the *Saccharomyces* and *Penicillium* genera (4, 10, 20, 34–37). *Saccharomyces* yeast species have anthropological significance with industrial and biotechnological applications and have been established as suitable model systems for evolutionary, genetics, and medical studies (38–40). The genetic and phenotypic diversity in other yeast genera, and the many relevant key insights that could be obtained from studying them, however, remain largely overlooked (3), particularly for studying adaptation.

Here, we looked outside the *Saccharomyces* genus to understand additional mechanisms of adaptation/domestication related to anthropogenic environments in other yeast species. We assessed domestication signatures in *Lachancea cidri*, a haploid species with an unknown sexual cycle, recently isolated from a cider fermentative environment in France (strain CBS2950), and from tree samples in Australia and Patagonia (29, 30). Our fermentation analyses demonstrate specific traits in the domesticated *L. cidri* CBS2950 strain that appear to provide a fitness advantage under fermentative conditions compared to wild strains. Our results are in agreement with previous reports where yeasts adapted to fermentative conditions showed an increased ethanol tolerance and more efficient amino acid consumption (3, 27, 33, 34, 41–48). Interestingly, we found a differential volatile compounds profile in wine fermentation by CBS2950. The CBS2950-derived wine has a less complex profile than that from native strains, a feature previously reported as a consequence of human selection (1, 4, 10, 39, 43, 49–51).

Using long– and short-read sequencing strategies, we demonstrated an enrichment of SVs in the CBS2950 strain compared to wild strains, which may partly be responsible for some of the phenotypic differences observed. Reports on other non-conventional yeasts have highlighted the presence of SVs with major impacts on the adaptation to new environments (8, 19, 24, 25), representing a rapid and efficient strategy to promote phenotypic changes to artificial environments. Indeed, several genomic modifications that impact the ability to ferment lactose and, thus, to adapt efficiently to dairy farmers’ environments were recently identified in *Kluyveromyces lactis* var. *lactis* (25). Moreover, *K. lactis* gained the ability to metabolize lactose from a horizontal gene transfer event from *Kluyveromyces marxianus* (19), emerging as a correlated response of the adaptation to the dairy processes (19, 25). Similarly, the acquisition of a cluster of *GAL* genes and the expansion and functional diversification of *MAL* genes were reported as domestication signatures to dairy and bread products in *Torulaspora delbrueckii* (8). Similar observations have been made in plants and fish, such as the Asian rice (*Oryza sativa*) and lake whitefish (*Coregonus* sp.), SVs contribute substantially to reshaping the genome architecture, underlying speciation events, and species differentiation with unique traits to adapt to anthropogenic ecosystems (52, 53).

For *L. cidri*, its haploid condition might promote major genomic rearrangements to generate a cost-effective adaptive change. Although single-nucleotide polymorphisms were initially hypothesized to underlie most selectable variation (12), we demonstrate here that genetic rearrangements like SVs might represent a major source of genetic variation under fermentative conditions in this species. Chromosomal rearrangements can underlie adaptation by affecting the expression of genes located in the proximity of the SVs breakpoints via gain, loss, or movement of regulatory regions (54). Here, we report the effect of a large translocation on the *PUT4* gene, located in the SV breakpoint neighborhood. RNA-seq analyses showed higher expression levels of *PUT4* in the cider strain, suggesting that the SV shed light on the molecular origin of the phenotypic differences between strains. Interestingly, these higher expression levels agree with increased ethanol tolerance in the “domesticated” cider strain. Consistent with this, previous studies have shown that an increase in proline consumption greatly improves tolerance to high ethanol concentrations (33). Thus, the SV breakpoint may have played a role by improving ethanol tolerance, amino acid consumption, and microbial growth in CBS2950. SVs can influence gene expression and impair gene function, which may result in signatures of local adaptation (9).

Another common domestication signature that has been associated to SVs is SO_2_ tolerance (24, 34). While we did observe increased tolerance to SO_2_ in CBS2950 compared to wild strains, we did not detect an SV associated with the *SSU1* gene or in its regulatory region. The *SSU1* gene is located approximately 540 kb upstream of the LACI0G translocation, suggesting an unknown molecular mechanism by which the “domesticated” strain performs better under high SO_2_ concentrations. This mechanism may differ from that found in other domesticated *S. cerevisiae* strains.

In conclusion, we found that SVs may account for most of the genetic variation between cider and wild strains in *L. cidri*, with a stronger signal over SNPs. Although SVs represent a rare event in natural populations (55), they could allow organisms to adapt to anthropogenic habitats. In particular, such events may have resulted in the ability of *L. cidri* to efficiently consume amino acids and tolerate high ethanol concentrations, allowing it to thrive in a fermentative environment. The different “domestication signatures” we observed for *L. cidri* under fermentation are similar to those found in *S. cerevisiae* strains under similar conditions (ethanol tolerance, efficient consumption of amino acids, and SO_2_ tolerance), suggesting convergent adaptive changes to anthropogenic environments in yeasts, despite over 100 MYA of divergence. These findings provide additional insights into microbial domestication and broaden the perspective of the fitness effects of SVs in yeast species associated to anthropogenic niches.

## Materials and Methods

### Yeast strains

Nine *L. cidri* strains, representative of the different lineages in the species, were selected for long-read sequencing and genome assembly (Table S1, (29, 30)). The genome assembly and annotation of the *L. cidri* CBS2950 strain were obtained from the GRYC (Genome Resources for Yeast Chromosomes, INRA, France).

### Fermentation performance in wine and cider

Fermentation assays were performed under three different must conditions: Cider, Chardonnay, and Synthetic wine must (SWM). Cider fermentation was performed in natural green apple juice (Afe®). Chardonnay grape juice was obtained from Blewitt Springs, Clarendon Hills (South Australia). Chardonnay juice contained 195 g/L of sugar (equal amounts of glucose and fructose), 8 mg/L of free sulfur dioxide, and had a pH 3.31. Yeast assimilable nitrogen (YAN) was adjusted to 250 mg N/L by adding diammonium phosphate (DAP). SWM was prepared at 300 mg/ml YAN (SWM300) and 60 mg/ml YAN (SWM60) as previously described (56).

For each fermentation, the strains were initially grown under constant agitation in 10 mL of YPD medium (1% yeast extract, 2% peptone, and 2% glucose) for 24 h at 20 °C. 1 x 10^6^ cells/mL were inoculated into 50 mL of each must (in 200-mL flasks) and incubated at 25 °C with constant agitation. Cider and SWM fermentations were weighed every day to calculate the accumulated CO_2_ loss. Chardonnay fermentation samples were taken regularly to monitor fermentation by measuring sugar concentration in culture supernatants using high-performance liquid chromatography (HPLC). At the end of the fermentation, samples were centrifuged at 9000 x g for 10 min, and the supernatant was collected for extracellular metabolite determination as previously described (56). Volatile compounds were measured as previously described (18, 49).

### DNA extraction and long-read sequencing

DNA was extracted from native *L. cidri* strains as previously described by (57). Libraries were sequenced on an R9.4 flow cell using a Minion (Oxford Nanopore Technologies, UK). The raw fast5 files were transformed to fastq files and debarcoded using Guppy v5.0.14 with the “super high accuracy” model (https://nanoporetech.com/accuracy). Barcodes and adapters sequences were trimmed using Porechop (https://github.com/rrwick/Porechop) and filtered with Filtlong (https://github.com/rrwick/Filtlong) using a Phred score of 30. In addition, we used publicly available paired-end Illumina sequence data for each strain (30).

### Genome assembly, annotation, and SV detection

Genome assembly was performed with Fly v2.9 (--nano-hq –g 400m) (58). Additionally, three rounds of nanopolish (https://github.com/jts/nanopolish) and pilon (https://github.com/broad institute/pilon) were carried out. The raw assembly was polished using Illumina reads filtered with a Phred score of 30 (Burrows-Wheeler Aligner). The genome assembly was annotated using the LRSDAY pipeline (59) and the *L. cidri* CBS2950 reference genome as a model. To identify the SVs in *L. cidri* strains, we performed pairwise comparisons between the *de novo* long-read assemblies of the nine strains using MUM&Co v3.8 (60). The pipeline used MUMmer v4 (61) to perform whole-genome alignments and detect SVs ≥ 50 bp.

### Phylogenetic analysis

A maximum-likelihood phylogenetic tree was constructed using protein sequences predicted from *L. cidri* strains and published genomes of *the Lachancea* genus strains available in GRYC: *Lachancea mirantina*, *Lachancea kluyveri* and *Lachancea fermentati*. Ortholog-Finder v2.4.1 (62) was employed to identify orthologous protein groups among these *Lachancea* species. Subsequently, a total of 3,408 single-copy orthologs identified in all species were aligned with Muscle v3.8.15, after which poorly aligned positions were trimmed with Gblocks v0.91v. 3,408 alignments were concatenated to produce a maximum-likelihood tree with RAxML v8.2.12 (-f a –x 12345 –p 12345 –# 100 –m PROTGAMMAJTT –k).

### RNA sequencing and differential expression analysis

Gene expression analysis was performed on strains *L. cidri* CBS2950 and *L. cidri* LC1, which exhibited a significant difference in fermentative kinetics under low nitrogen conditions (SWM60). The two strains were fermented in SWM60 in triplicates and RNA was obtained as previously described (56). RNA integrity was confirmed using Fragment Analyzer (Agilent). RNA-seq libraries and reads trimming were performed as previously described by (56). Cleaned reads were mapped to the *L. cidri* CBS2950 genome assembly using HISAT2 v2.2.1 (––max-intronlen 25000 ––dta-cufflinks) (63). Differentially expressed gene (DEG) analysis was performed using the DESeq2 package in R v4.1.2, comparing the two strains (64). Genes with a FDR < 0.05 and an absolute value of fold change > 1.5 were considered DEGs for the comparison, *L. cidri* CBS2950 vs. *L. cidri* LC1. Gene Ontology analysis was performed with the R package enrichGO.

### Growth under proline as nitrogen source

Cells were pre-cultivated at 20 °C without agitation for 48 h in 96-well plates containing 200-μL YNB (Yeast Nitrogen Base w/o ammonium and amino acids; Difco) medium supplemented with 2% (w/v) glucose. A volume of 10 μl of pre-inoculum was used to inoculate a new 96-well plate containing 200 μl of YNB w/o ammonium and amino acids with different concentrations of proline (200, 100, and 50 mg mL YAN) to an optical density (OD_600_) of 0.03–0.1. OD_600_ for each well was measured at 620 nm every 30 min for 250 h. As a control, we used a combination of four different amino acids (Glutamic acid, Aspartic acid, Alanine, and Leucine) at 200 and 50 mg/mL YAN. From these data, three parameters were estimated: lag phase, growth rate (μmax), and maximum OD using the GrowthRates software (65).

## Ethanol and sulfite tolerance screening

Ethanol tolerance was evaluated in agar plates (YNB-agar 2%) supplemented with different ethanol concentrations (6, 8, 12, and 15 % v/v). Yeast cells were cultivated to exponential phase, then spotted on 10-fold serial dilutions on agar plates and incubated at 20 °C for 3 days. Sulfite tolerance assays were performed as described previously (24). Microplates were sealed with Breathe-Easy® sealing membranes (Diversified Biotech, USA) and incubated at 28 °C. OD_600_ was measured after five days using a Tecan infinite M200 plate reader (Tecan, Austria).

## Statistical analyses

All phenotype experiments were carried out at least in triplicate. All statistical analyses (T-test, one-way ANOVA with correction for multiple comparisons) were performed in R software (66). A *p*-value < 0.05 was considered statistically significant.

## Data availability

All fastq sequences were deposited in the National Center for Biotechnology Information (NCBI) as a Sequence Read Archive under the BioProject accession number PRJNA978517 (www.ncbi.nlm.nih.gov/bioproject/978517).

## Acknowledgments

This research was funded by Agencia Nacional de Investigación y Desarrollo (ANID) FONDECYT (1220026) and ANID – Programa Iniciativa Científica Milenio ICN17_022 and NCN2021_050. PV is supported by ANID FONDECYT POSTDOCTORADO [grant 3200575]. RN is supported by FONDECYT grant [1180917] and NCN2021_050. This research was supported by the supercomputing infrastructure of the NLHPC (ECM-02). FC, RN, PV, TP, SO, and GF were funded by PROGRAMA DE COOPERACIÓN CIENTÍFICA ECOS-CONICYT ECOS180003. We acknowledge Fundación Ciencia & Vida for providing infrastructure, laboratory space, and equipment for experiments. We thank Alejandro Montenegro-Montero for language and editorial support.

## Author Contributions

**Conceptualization:** P.V., F.A.C. **Formal analysis:** P.V. **Investigation:** P.V., S.O., N.A., F.M.G., J.BP., K.U., T.P., M.S, F.A.C. **Writing – original draft:** P.V., F.A.C.

**Writing – review & editing:** P.V., R.F.N., C.V., G.F., F.A.C.

## Expanded View Figures legends and tables

**Figure S1.**
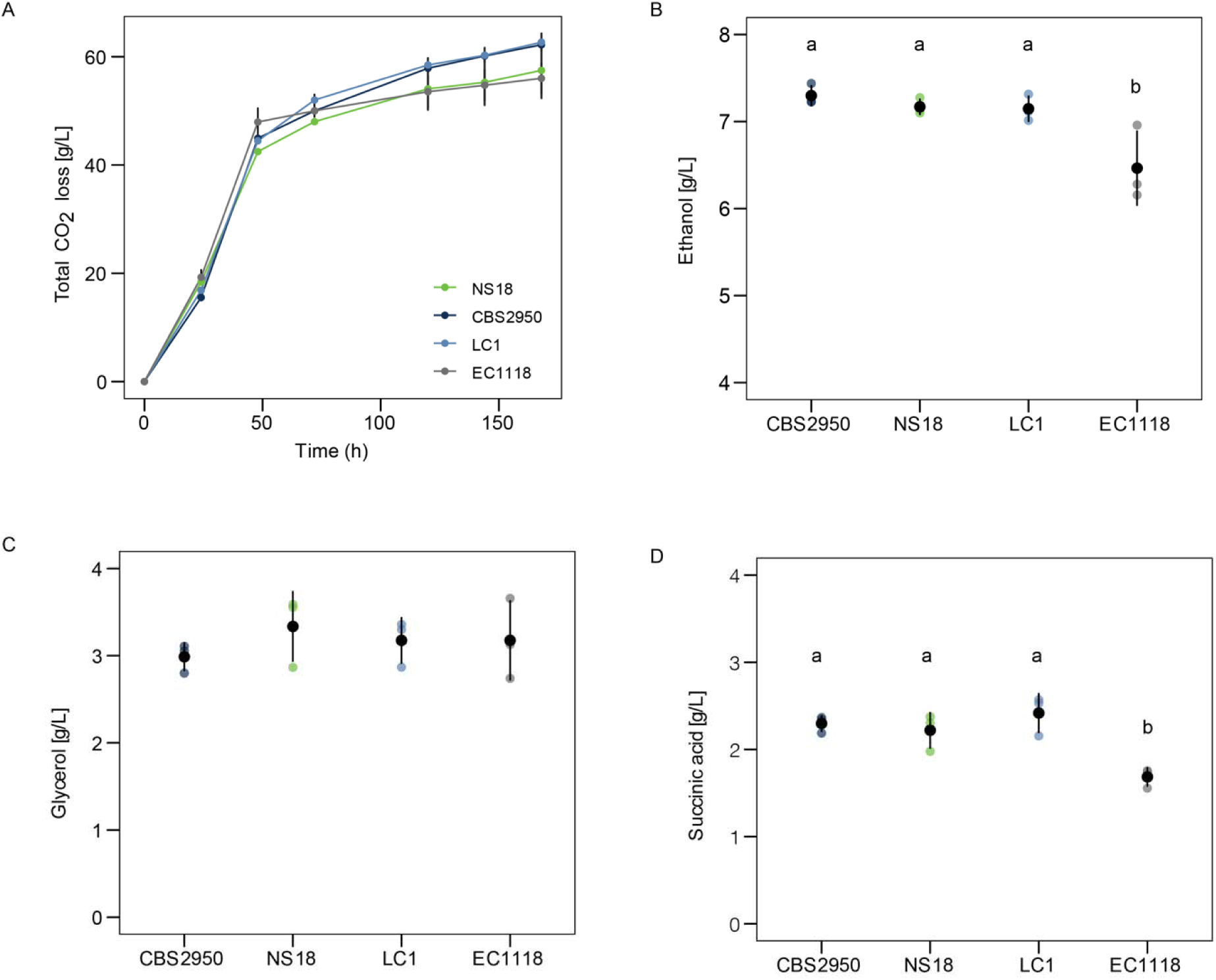
*Lachancea cidri* fermentation performance in Cider. (A) Fermentation kinetics of wild and human-related strains. (B) Ethanol production [g/L]. (C) Glycerol production [g/L]. (D) Succinic acid production [g/L]. Different letters reflect statistically differences between strains with a *p*-value < 0.05, one-way analysis of variance (ANOVA).

**Figure S2.**
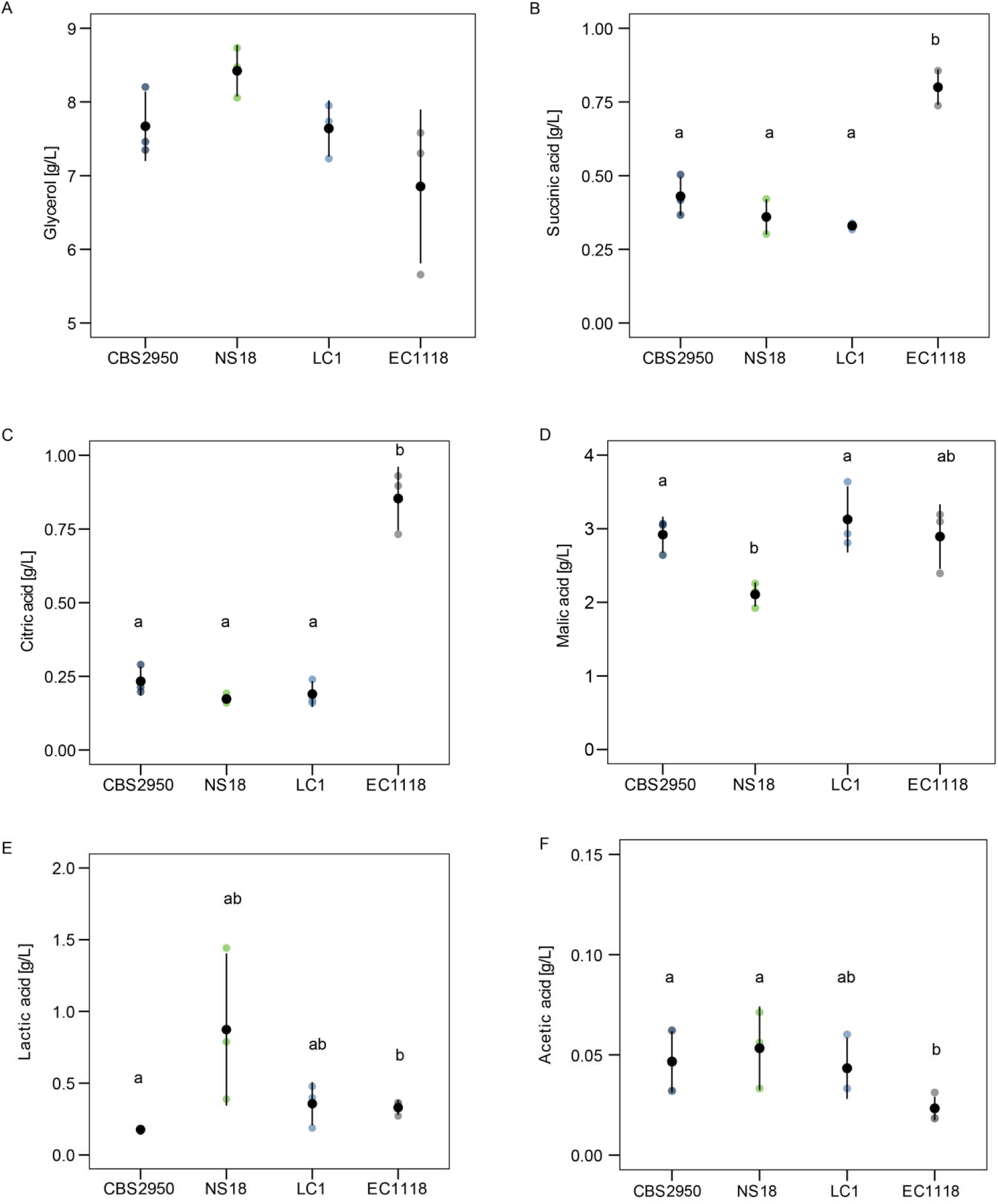
Metabolites produced at the end of Synthetic Wine Must fermentation. (A). Glycerol production [g/L]. (B) Succinic acid production [g/L] (C) Citric acid production [g/L]. (D) Malic acid production [g/L]. (E) Lactic acid production [g/L]. (F) Acetic acid production [g/L]. Different letters reflect statistically differences between strains with a *p*-value < 0.05, one-way analysis of variance (ANOVA).

**Figure S3.**
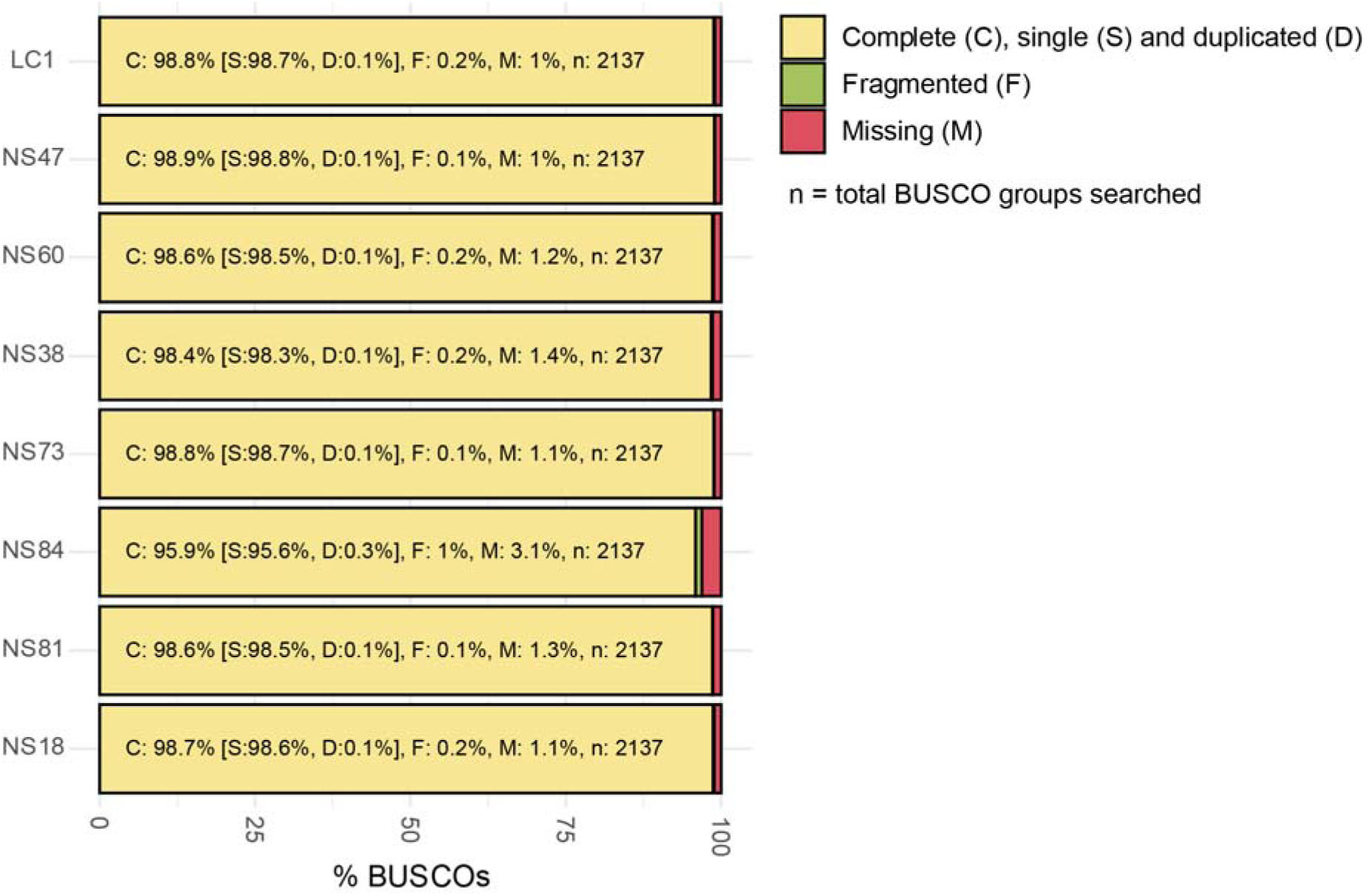
BUSCO’s completeness of each assembly.

**Figure S4.**
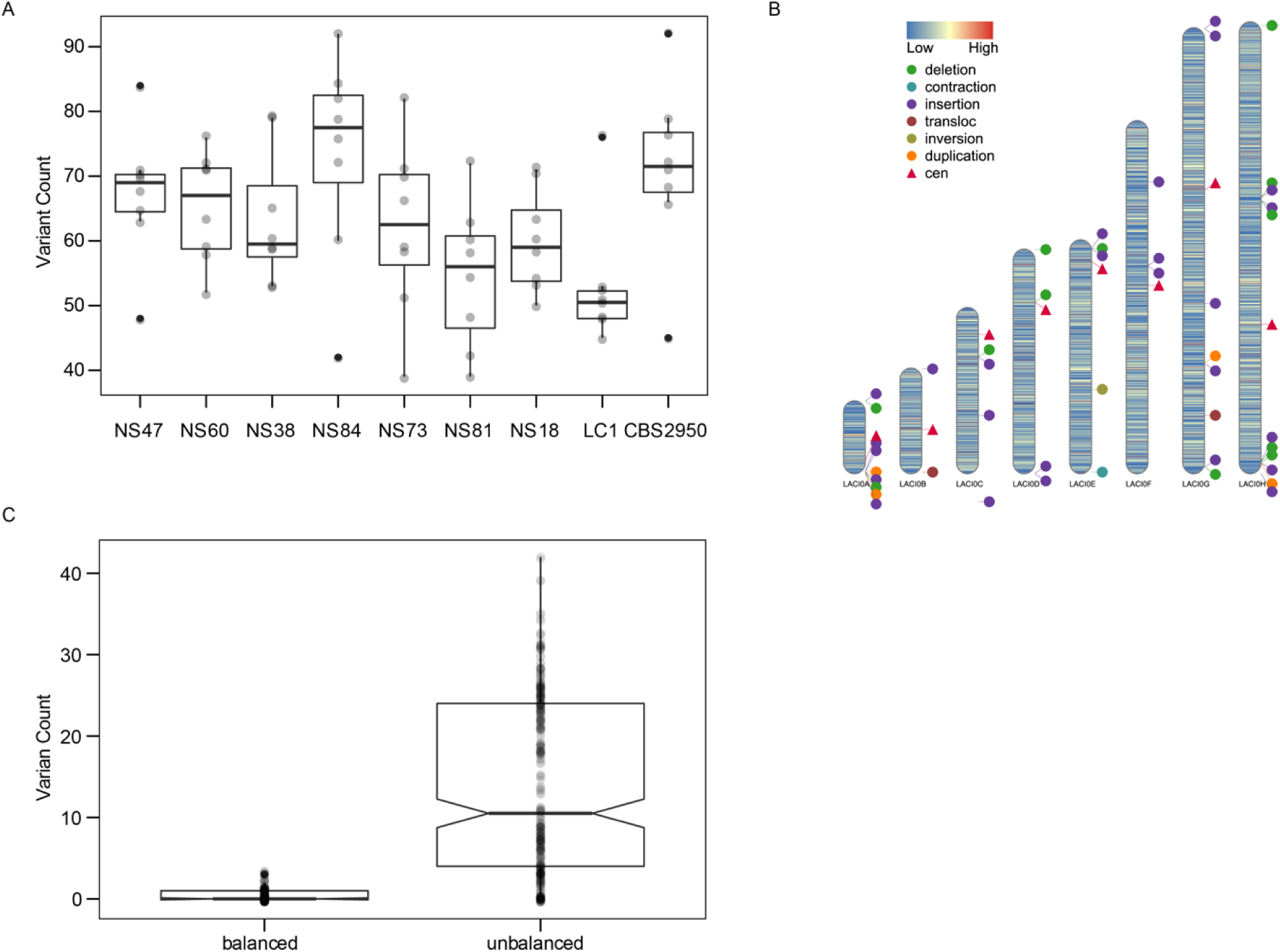
Structural variations within *Lachancea cidri* strains. (A). The range of total structural variation counts found for each genome serves as the reference genome. (B). SV distribution in the genome between LC1 and CBS2950 (C). Balanced and Unbalanced variant count.

**Figure S5.**
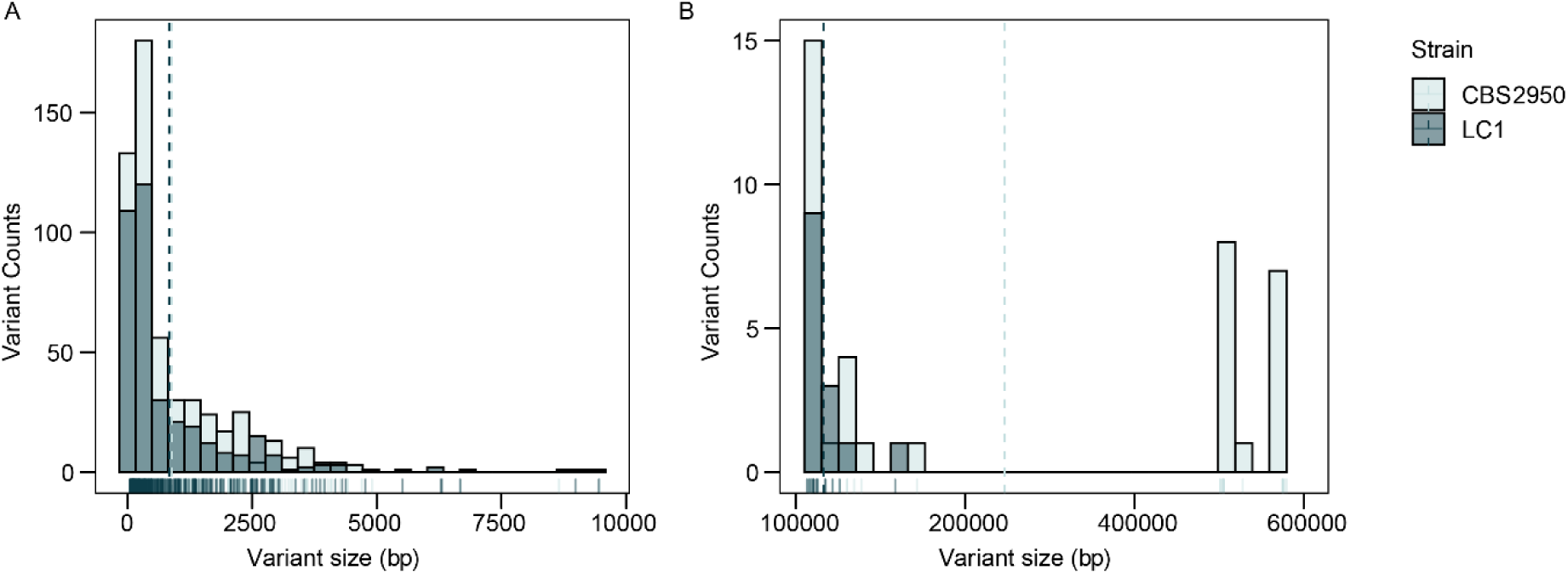
Distribution of SV sizes between *L. cidri* CBS2950 and LC1. (A). SV < 10,000 bp. (B). SV > 10,000 bp.

**Figure S6.**
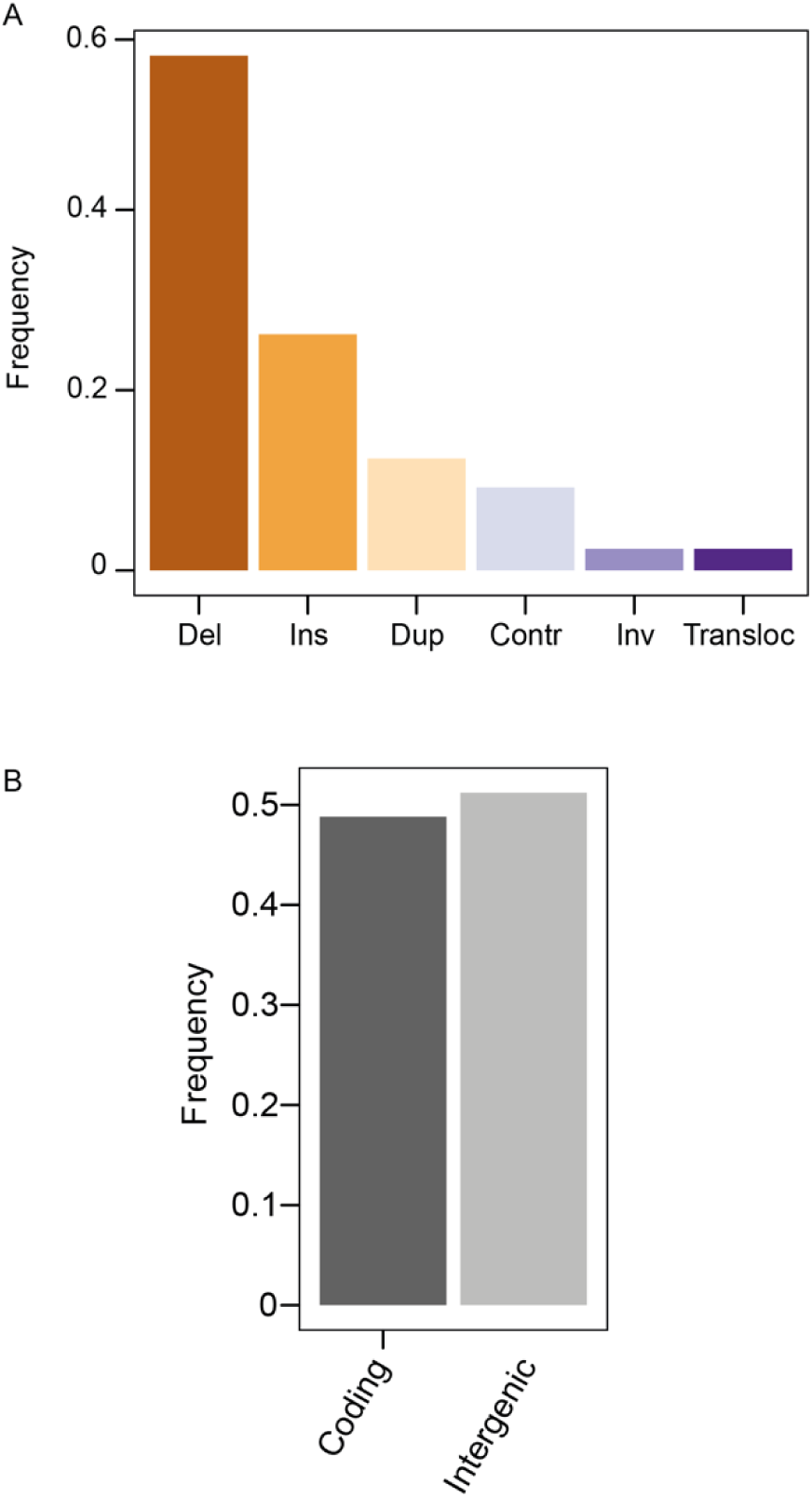
(A) Frequency of SVs between CBS2950 and LC1 (Del for deletions, Ins for insertions, Dup for duplications, Inv for inversions, Contr for contractions, and Transloc for translocations) (B) Frequency of SVs located in coding or non-coding regions between CBS2950 and LC1 strains.

**Figure S7.**
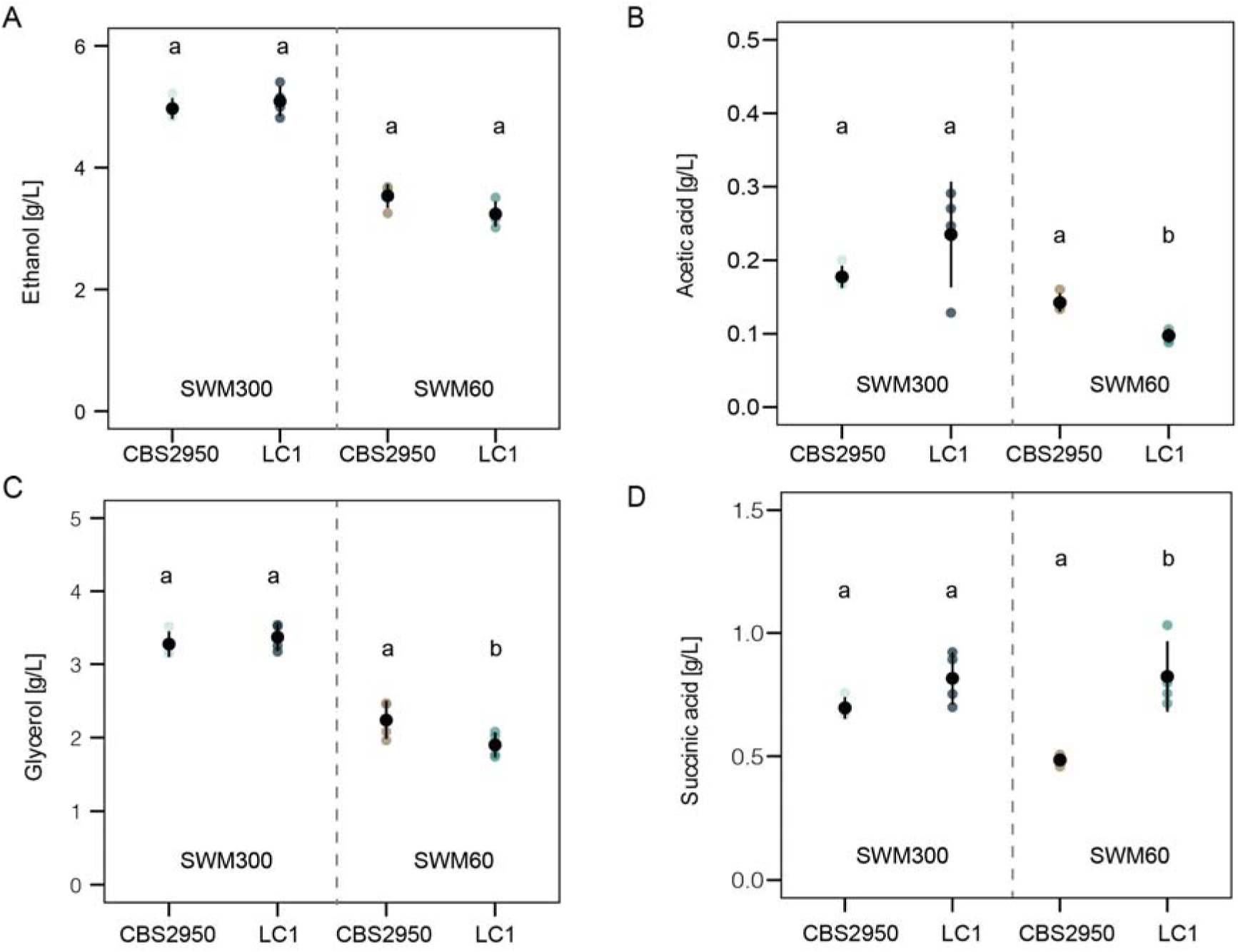
Metabolites produced at the end of Synthetic Wine Must fermentation. (A). Ethanol [g/L]. (B). Acetic acid [g/L] (C). Glycerol [g/L]. (D) Succinic acid [g/L]. Different letters reflect statistically differences between strains with a *p*-value < 0.05, one-way analysis of variance (ANOVA).

**Figure S8.**
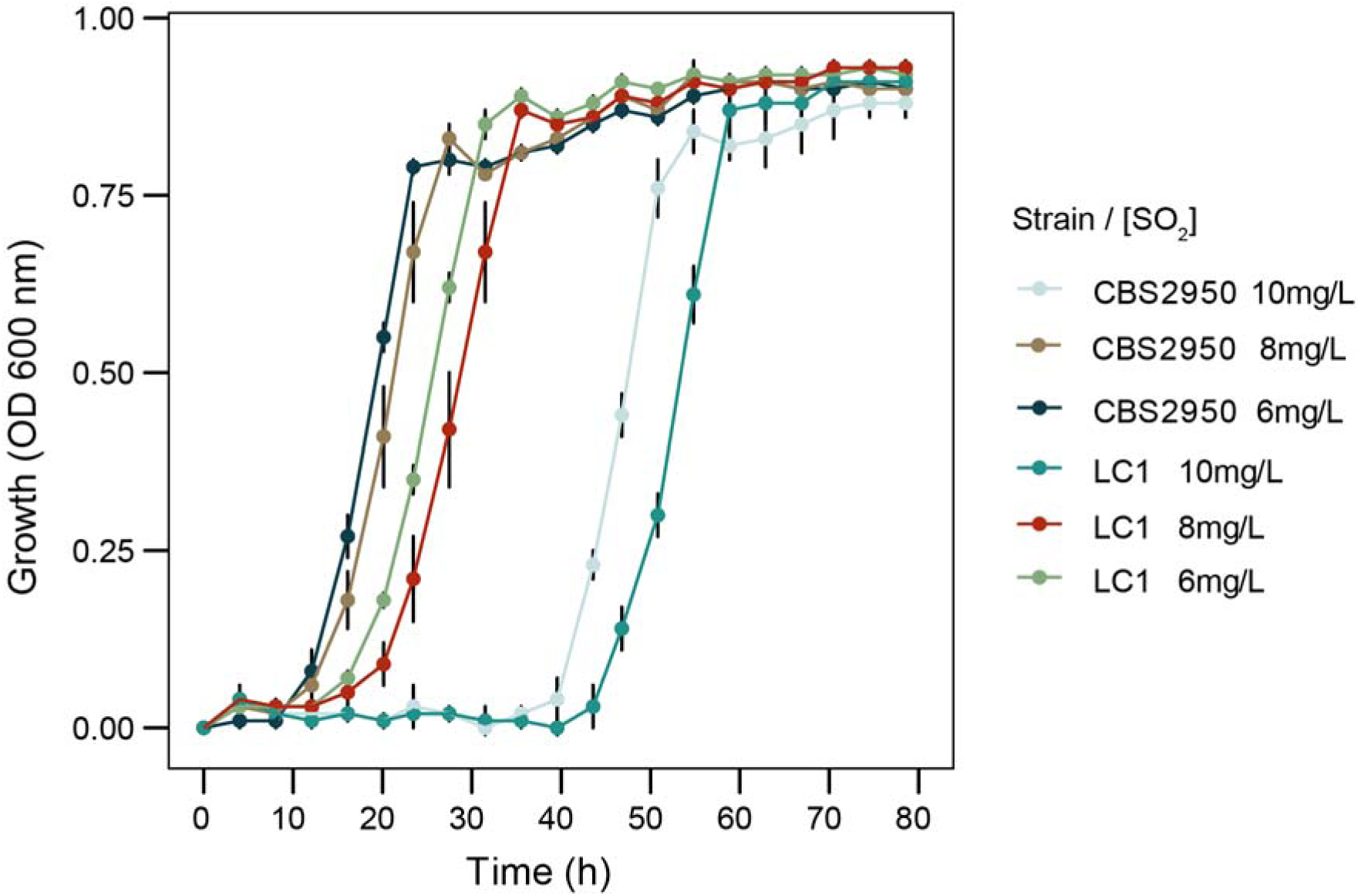
Growth kinetics at different sulfite concentrations.

## References

1. Purugganan MD. 2022. What is domestication? Trends Ecol Evol 37:663–671.

2. Larson G, Piperno DR, Allaby RG, Purugganan MD, Andersson L, Arroyo-Kalin M, Barton L, Climer Vigueira C, Denham T, Dobney K, Doust AN, Gepts P, Gilbert MT, Gremillion KJ, Lucas L, Lukens L, Marshall FB, Olsen KM, Pires JC, Richerson PJ, Rubio de Casas R, Sanjur OI, Thomas MG, Fuller DQ. 2014. Current perspectives and the future of domestication studies. Proc Natl Acad Sci U S A 111:6139–46.

3. Steensels J, Gallone B, Voordeckers K, Verstrepen KJ. 2019. Domestication of Industrial Microbes. Curr Biol 29:R381–R393.

4. Gibbons JG, Rinker DC. 2015. The genomics of microbial domestication in the fermented food environment. Curr Opin Genet Dev 35:1–8.

5. Liti G, Schacherer J. 2011. The rise of yeast population genomics. C R Biol 334:612–9.

6. Peter J, Schacherer J. 2016. Population genomics of yeasts: towards a comprehensive view across a broad evolutionary scale. Yeast 33:73–81.

7. Steenwyk JL, Rokas A. 2018. Copy Number Variation in Fungi and Its Implications for Wine Yeast Genetic Diversity and Adaptation. Front Microbiol 9:288.

8. Silva M, Pontes A, Franco-Duarte R, Soares P, Sampaio JP, Sousa MJ, Brito PH. 2022. A glimpse at an early stage of microbe domestication revealed in the variable genome of Torulaspora delbrueckii, an emergent industrial yeast. Mol Ecol doi:10.1111/mec.16428.

9. Hamala T, Wafula EK, Guiltinan MJ, Ralph PE, dePamphilis CW, Tiffin P. 2021. Genomic structural variants constrain and facilitate adaptation in natural populations of Theobroma cacao, the chocolate tree. Proc Natl Acad Sci U S A 118.

10. Ina Bodinaku JS, Allison B. Connors, Jacob L. Steenwyk, Megan N. Biango-Daniels, Erik K. Kastman, Antonis Rokas, Albert Robbat, Benjamin E. Wolfea. 2019. Rapid Phenotypic and Metabolomic Domestication of Wild Penicillium Molds on Cheese. mBio 10.

11. Steenwyk J, Rokas A. 2017. Extensive Copy Number Variation in Fermentation-Related Genes Among Saccharomyces cerevisiae Wine Strains. G3 (Bethesda) 7:1475–1485.

12. Wellenreuther M, Merot C, Berdan E, Bernatchez L. 2019. Going beyond SNPs: The role of structural genomic variants in adaptive evolution and species diversification. Mol Ecol 28:1203–1209.

13. García-Ríos E, Nuévalos M, Barrio E, Puig S, Guillamón JM. 2019. A new chromosomal rearrangement improves the adaptation of wine yeasts to sulfite. Environmental Microbiology 21:1771–1781.

14. Gorkovskiy A, Verstrepen KJ. 2021. The Role of Structural Variation in Adaptation and Evolution of Yeast and Other Fungi. Genes (Basel) 12.

15. Wong VL, Ellison CE, Eisen MB, Pachter L, Brem RB. 2014. Structural Variation among Wild and Industrial Strains of Penicillium chrysogenum. PLOS ONE 9:e96784.

16. Cheeseman K, Ropars J, Renault P, Dupont J, Gouzy J, Branca A, Abraham AL, Ceppi M, Conseiller E, Debuchy R, Malagnac F, Goarin A, Silar P, Lacoste S, Sallet E, Bensimon A, Giraud T, Brygoo Y. 2014. Multiple recent horizontal transfers of a large genomic region in cheese making fungi. Nat Commun 5:2876.

17. Macias LG, Flores MG, Adam AC, Rodriguez ME, Querol A, Barrio E, Lopes CA, Perez-Torrado R. 2021. Convergent adaptation of Saccharomyces uvarum to sulfite, an antimicrobial preservative widely used in human-driven fermentations. PLoS Genet 17:e1009872.

18. Varela C, Barker A, Tran T, Borneman A, Curtin C. 2017. Sensory profile and volatile aroma composition of reduced alcohol Merlot wines fermented with Metschnikowia pulcherrima and Saccharomyces uvarum. Int J Food Microbiol 252:1–9.

19. Varela JA, Puricelli M, Ortiz-Merino RA, Giacomobono R, Braun-Galleani S, Wolfe KH, Morrissey JP. 2019. Origin of Lactose Fermentation in Kluyveromyces lactis by Interspecies Transfer of a Neo-functionalized Gene Cluster during Domestication. Curr Biol 29:4284–4290 e2.

20. Zimmer A, Durand C, Loira N, Durrens P, Sherman DJ, Marullo P. 2014. QTL dissection of Lag phase in wine fermentation reveals a new translocation responsible for Saccharomyces cerevisiae adaptation to sulfite. PLoS One 9:e86298.

21. Fleiss A, O’Donnell S, Fournier T, Lu W, Agier N, Delmas S, Schacherer J, Fischer G. 2019. Reshuffling yeast chromosomes with CRISPR/Cas9. PLoS Genet 15:e1008332.

22. Perez-Ortin JE, Querol A, Puig S, Barrio E. 2002. Molecular characterization of a chromosomal rearrangement involved in the adaptive evolution of yeast strains. Genome Res 12:1533–9.

23. Cubillos FA, Gibson B, Grijalva-Vallejos N, Krogerus K, Nikulin J. 2019. Bioprospecting for brewers: Exploiting natural diversity for naturally diverse beers. Yeast 36:383–398.

24. Varela C, Bartel C, Roach M, Borneman A, Curtin C. 2019. Brettanomyces bruxellensis SSU1 Haplotypes Confer Different Levels of Sulfite Tolerance When Expressed in a Saccharomyces cerevisiae SSU1 Null Mutant. Appl Environ Microbiol 85.

25. Friedrich A, Gounot J-S, Tsouris A, Bleykasten C, Freel K, Caradec C, Schacherer J. 2023. Contrasting genomic evolution between domesticated and wild Kluyveromyces lactis yeast populations. Genome Biology and Evolution doi:10.1093/gbe/evad004:evad004.

26. Albertin W, Chasseriaud L, Comte G, Panfili A, Delcamp A, Salin F, Marullo P, Bely M. 2014. Winemaking and bioprocesses strongly shaped the genetic diversity of the ubiquitous yeast Torulaspora delbrueckii. PLoS One 9:e94246.

27. Valles BS, Bedrinana RP, Tascon NF, Simon AQ, Madrera RR. 2007. Yeast species associated with the spontaneous fermentation of cider. Food Microbiol 24:25–31.

28. Coton E, Coton M, Levert D, Casaregola S, Sohier D. 2006. Yeast ecology in French cider and black olive natural fermentations. Int J Food Microbiol 108:130–5.

29. Vakirlis N, Sarilar V, Drillon G, Fleiss A, Agier N, Meyniel JP, Blanpain L, Carbone A, Devillers H, Dubois K, Gillet-Markowska A, Graziani S, Huu-Vang N, Poirel M, Reisser C, Schott J, Schacherer J, Lafontaine I, Llorente B, Neuveglise C, Fischer G. 2016. Reconstruction of ancestral chromosome architecture and gene repertoire reveals principles of genome evolution in a model yeast genus. Genome Res 26:918–32.

30. Villarreal P, Villarroel CA, O’Donnell S, Agier N, Quintero-Galvis JF, Pena TA, Nespolo RF, Fischer G, Varela C, Cubillos FA. 2022. Late Pleistocene-dated divergence between South Hemisphere populations of the non-conventional yeast L. cidri. Environ Microbiol doi:10.1111/1462-2920.16103.

31. Varela C, Sundstrom J, Cuijvers K, Jiranek V, Borneman A. 2020. Discovering the indigenous microbial communities associated with the natural fermentation of sap from the cider gum Eucalyptus gunnii. Sci Rep 10:14716.

32. Kurtzman C. 2003. Phylogenetic circumscription of, and other members of the Saccharomycetaceae, and the proposal of the new genera, and. FEMS Yeast Research 4:233–245.

33. Takagi H, Taguchi J, Kaino T. 2016. Proline accumulation protects Saccharomyces cerevisiae cells in stationary phase from ethanol stress by reducing reactive oxygen species levels. Yeast 33:355–63.

34. Garcia-Rios E, Nuevalos M, Barrio E, Puig S, Guillamon JM. 2019. A new chromosomal rearrangement improves the adaptation of wine yeasts to sulfite. Environ Microbiol 21:1771–1781.

35. Peter J, De Chiara M, Friedrich A, Yue JX, Pflieger D, Bergstrom A, Sigwalt A, Barre B, Freel K, Llored A, Cruaud C, Labadie K, Aury JM, Istace B, Lebrigand K, Barbry P, Engelen S, Lemainque A, Wincker P, Liti G, Schacherer J. 2018. Genome evolution across 1,011 Saccharomyces cerevisiae isolates. Nature 556:339–344.

36. Lage P, Sampaio-Marques B, Ludovico P, Mira NP, Mendes-Ferreira A. 2019. Transcriptomic and chemogenomic analyses unveil the essential role of Com2-regulon in response and tolerance of Saccharomyces cerevisiae to stress induced by sulfur dioxide. Microb Cell 6:509–523.

37. Pitt JI, Cruickshank RH, Leistner L. 1986. Penicillium commune, P. camembertii, the origin of white cheese moulds, and the production of cyclopiazonic acid. Food Microbiology 3:363–371.

38. Borneman AR, Pretorius IS. 2015. Genomic insights into the Saccharomyces sensu stricto complex. Genetics 199:281–91.

39. Gibson B, Geertman JA, Hittinger CT, Krogerus K, Libkind D, Louis EJ, Magalhaes F, Sampaio JP. 2017. New yeasts-new brews: modern approaches to brewing yeast design and development. FEMS Yeast Res 17.

40. Cubillos FA. 2016. Exploiting budding yeast natural variation for industrial processes. Curr Genet 62:745–751.

41. Garcia-Rios E, Guillen A, de la Cerda R, Perez-Traves L, Querol A, Guillamon JM. 2018. Improving the Cryotolerance of Wine Yeast by Interspecific Hybridization in the Genus Saccharomyces. Front Microbiol 9:3232.

42. Maicas S. 2020. The Role of Yeasts in Fermentation Processes. Microorganisms 8.

43. Minebois R, Lairon-Peris M, Barrio E, Perez-Torrado R, Querol A. 2021. Metabolic differences between a wild and a wine strain of Saccharomyces cerevisiae during fermentation unveiled by multi-omic analysis. Environ Microbiol 23:3059–3076.

44. Origone AC, Del Monaco SM, Avila JR, Gonzalez Flores M, Rodriguez ME, Lopes CA. 2017. Tolerance to winemaking stress conditions of Patagonian strains of Saccharomyces eubayanus and Saccharomyces uvarum. J Appl Microbiol 123:450–463.

45. Padilla B, Gil JV, Manzanares P. 2016. Past and Future of Non-Saccharomyces Yeasts: From Spoilage Microorganisms to Biotechnological Tools for Improving Wine Aroma Complexity. Front Microbiol 7:411.

46. Shiroma S, Jayakody LN, Horie K, Okamoto K, Kitagaki H. 2014. Enhancement of ethanol fermentation in Saccharomyces cerevisiae sake yeast by disrupting mitophagy function. Appl Environ Microbiol 80:1002–12.

47. Varela C. 2016. The impact of non-Saccharomyces yeasts in the production of alcoholic beverages. Appl Microbiol Biotechnol 100:9861–9874.

48. Su Y, Seguinot P, Sanchez I, Ortiz-Julien A, Heras JM, Querol A, Camarasa C, Guillamon JM. 2020. Nitrogen sources preferences of non-Saccharomyces yeasts to sustain growth and fermentation under winemaking conditions. Food Microbiol 85:103287.

49. Urbina K, Villarreal P, Nespolo RF, Salazar R, Santander R, Cubillos FA. 2020. Volatile Compound Screening Using HS-SPME-GC/MS on Saccharomyces eubayanus Strains under Low-Temperature Pilsner Wort Fermentation. Microorganisms 8.

50. Gibson B, Dahabieh M, Krogerus K, Jouhten P, Magalhaes F, Pereira R, Siewers V, Vidgren V. 2020. Adaptive Laboratory Evolution of Ale and Lager Yeasts for Improved Brewing Efficiency and Beer Quality. Annu Rev Food Sci Technol 11:23–44.

51. Krogerus K, Arvas M, De Chiara M, Magalhaes F, Mattinen L, Oja M, Vidgren V, Yue JX, Liti G, Gibson B. 2016. Ploidy influences the functional attributes of de novo lager yeast hybrids. Appl Microbiol Biotechnol 100:7203–22.

52. Merot C, Stenlokk KSR, Venney C, Laporte M, Moser M, Normandeau E, Arnyasi M, Kent M, Rougeux C, Flynn JM, Lien S, Bernatchez L. 2022. Genome assembly, structural variants, and genetic differentiation between lake whitefish young species pairs (Coregonus sp.) with long and short reads. Mol Ecol doi:10.1111/mec.16468.

53. Yixuan Kou, Yi Liao, Tuomas Toivainen, Yuanda Lv, Xinmin Tian, J.J. Emerson, Brandon S. Gaut aYZ. 2020. Evolutionary Genomics of Structural Variation in Asian Rice (Oryza sativa) Domestication. Mol Biol Evol 37(1):3507–3524.

54. Roesti M, Gilbert KJ, Samuk K. 2022. doi:10.1101/2022.05.02.490344.

55. O’Donnell S, Yue J-X, Saada OA, Agier N, Caradec C, Cokelaer T, Chiara MD, Delmas S, Dutreux F, Fournier T, Friedrich A, Kornobis E, Li J, Miao Z, Tattini L, Schacherer J, Liti G, Fischer G. 2022. 142 telomere-to-telomere assemblies reveal the genome structural landscape in Saccharomyces cerevisiae bioRxiv doi:10.1101/2022.10.04.510633.

56. Villarroel CA BM, Canessa P, Cubillos FA. 2021. Uncovering divergence in gene expression regulation in the adaptation of yeast to nitrogen scarcity. mSystems 6:e00466–21 6.

57. Nespolo RF, Villarroel CA, Oporto CI, Tapia SM, Vega-Macaya F, Urbina K, De Chiara M, Mozzachiodi S, Mikhalev E, Thompson D, Larrondo LF, Saenz-Agudelo P, Liti G, Cubillos FA. 2020. An Out-of-Patagonia migration explains the worldwide diversity and distribution of Saccharomyces eubayanus lineages. PLoS Genet 16:e1008777.

58. Kolmogorov M, Yuan J, Lin Y, Pevzner PA. 2019. Assembly of long, error-prone reads using repeat graphs. Nat Biotechnol 37:540–546.

59. Yue JX, Liti G. 2018. Long-read sequencing data analysis for yeasts. Nat Protoc 13:1213–1231.

60. O’Donnell S, Fischer G. 2020. MUM&Co: accurate detection of all SV types through whole-genome alignment. Bioinformatics 36:3242–3243.

61. Marcais G, Delcher AL, Phillippy AM, Coston R, Salzberg SL, Zimin A. 2018. MUMmer4: A fast and versatile genome alignment system. PLoS Comput Biol 14:e1005944.

62. Emms DM, Kelly S. 2019. OrthoFinder: phylogenetic orthology inference for comparative genomics. Genome Biol 20:238.

63. Kim D, Paggi JM, Park C, Bennett C, Salzberg SL. 2019. Graph-based genome alignment and genotyping with HISAT2 and HISAT-genotype. Nat Biotechnol 37:907–915.

64. Love MI, Huber W, Anders S. 2014. Moderated estimation of fold change and dispersion for RNA-seq data with DESeq2. Genome Biol 15:550.

65. Hall BG, Acar H, Nandipati A, Barlow M. 2014. Growth rates made easy. Mol Biol Evol 31:232–8.

66. Team TRC. 2014. R: A Language and Environment for Statist.

